# Structural insights into how metallochaperones UreE and UreG interact to deliver a toxic metal to urease

**DOI:** 10.64898/2025.12.08.692910

**Authors:** Chun-Long Chan, Longson Tsz Hin Pang, Tung Choi, Ka-Chun Chan, Ka Lung Tsang, Ka Ming Lee, Kam-Bo Wong

## Abstract

Maturation of urease, a virulence factor for *Helicobacter pylori* infection, requires the delivery of nickel ions to the metalloenzyme. To avoid cytotoxicity, nickel ions are transferred within protein complexes of metallochaperones to ensure that the toxic metal do not escape into the cytoplasm. In the urease maturation pathway, metallochaperone UreG receives its nickel ions by forming a 2:2 complex with another metallochaperone UreE. Using C-terminal truncation variants of UreE [UreE(1-158) and UreE(1-148)], we determined the crystal structures of the UreE_2_G_2_ complex bound with nickel ions and GMPPNP, a non-hydrolysable GTP analogue. UreE-UreG interactions are asymmetric, where the UreG dimer mainly interacts with the proximal UreE. GTP binding induces conformational changes in the G2 and CPH motifs of distal UreG that enable its anchorage to the UreE_2_G_2_ complex. His68 of distal UreG moves towards and chelates a nickel ion at the UreE binding site. Formation of the UreE_2_G_2_ complex juxtaposes the nickel binding sites of UreE and UreG. Nickel transfer from UreE to UreG induces conformational changes that weaken UreE-UreG interactions, thereby facilitating UreG dissociation. We further demonstrated that the hydrogenase maturation factor HypA, providing the nickel source, could activate urease in vitro through protein-protein interaction with wild-type UreE but not with its truncation variants. UreG binding induces conformational changes in the C-terminal tail of UreE, promoting dissociation of HypA-UreE complex. Our work presents a paradigm on how GTP and nicking binding allosterically regulate the formation of a metallochaperone complex to facilitate nickel transfer.

**Significance:** Our work provides insights into how cells solve the problem of trafficking a toxic metal, nickel, to the active site of urease. Colonization of *Helicobacter pylori* in acidic human stomach requires the biosynthesis of active urease, which involves the delivery of the toxic nickel ions to the active site of the metalloenzyme. To avoid cytotoxicity, nickel ions are transported from one metallochaperone to another via the formation of protein complexes so that the toxic metal ions do not escape into the cytoplasm. Supported by structural and biochemical evidence, we present a paradigm on how GTP and nickel binding allosterically promote the formation of a complex between metallochaperones UreE and UreG to facilitate nickel transfer between the two proteins.

## Introduction

*Helicobacter pylori* infects half of the human population and is associated with an increased risk of peptic ulcers and gastric cancer (1). *H. pylori* is the only pathogen that can colonize the acidic human stomach because it produces large amount of urease, which hydrolyze urea into ammonia that helps the bacterium to maintain an intracellular neutral pH (2). The urease is a nickel (Ni) enzyme containing two Ni(II) ions in the active site (3, 4). The apo-urease is inactive, and biosynthesis of active urease requires a maturation process that deliver the nickel ions to the active site of urease (5).

Nickel ion is cytotoxic as it can displace weaker ions such as magnesium ions in the active site of essential enzymes like GTPases (6). As a result, the metal ion is tightly regulated so that there are essentially no free nickel ions in the bacterial cells (7, 8). The urease maturation pathway represents a paradigm on how cells solve the problem of delivering a toxic metal ion from one protein to another. To avoid cytotoxicity, cells have evolved metallochaperones, or metal carrier proteins, to transport Ni(II) ions (5). There are four urease accessory proteins that are involved in the nickel delivery to the urease: UreD (formerly called UreH), UreE, UreF and UreG (Fig. 1A). UreE is a homodimer that can bind a Ni(II) ion at the dimeric interface with a conserved GNRH motif (9–11). UreG is a SIMIBI (Signal recognition particle, MinD and BioD) GTPase that can undergo Ni/GTP-dependent dimerization that chelates a Ni(II) ion in a square-planar geometry with a conserved CPH motif at the dimeric interface (12, 13). UreF and UreD forms a UreFD complex that interacts GDP-bound UreG to form a 2:2:2 heterotrimeric UreGFD complex (12, 14). In the presence of GTP, UreG dissociates from the UreGFD complex and forms a UreE_2_G_2_ complex with UreE (13, 15) (Fig. 1A). After receiving its nickel ions from the UreE_2_G_2_ complex, UreG then forms an activation complex with UreFD and apo-urease. Upon GTP hydrolysis, the nickel ion is released from UreG, pass through a protein tunnel within the activation complex, and reach the active site of urease (16) (Fig. 1A).

**Figure 1.**
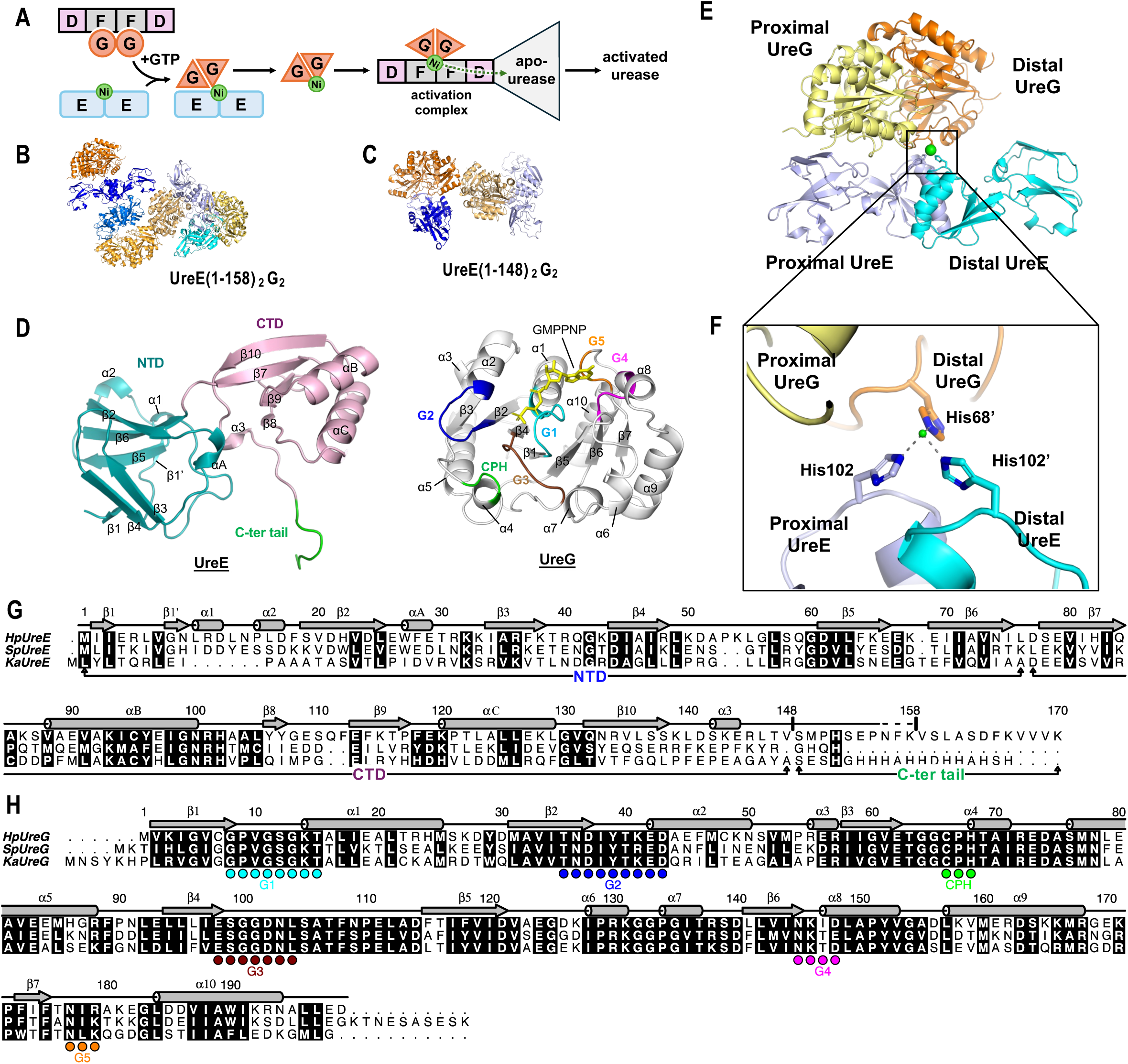
Crystal structures of UreE_2_G_2_ in complex with nickel and GMPPNP. (A) In the presence of GTP, UreG dissociates from the UreGFD complex and forms a 2:2 complex with UreE. UreG gets its nickel ion from the UreE_2_G_2_ complex and then forms an activation complex with UreFD and apo-urease. Upon GTP hydrolysis, the nickel ion is delivered to the active site of urease via a protein tunnel within the activation complex. (B-C) The structures of (B) UreE(1-158)_2_G_2_ and (C) UreE(1-148)_2_G_2_ in complex with Ni(II) and GMPPNP were determined at a resolution of 3.10 Å and 2.85 Å, respectively. In the asymmetry unit of UreE(1-158)_2_G_2_ and UreE(1-148)_2_G_2_, four and two copies of the UreE_2_G_2_ complexes were found, respectively. UreE and UreG dimers are in different shades of orange and blue, respectively. (D) Overall folds of UreE (left panel) and UreG (right panel). The C-terminal tail, N-terminal (NTD) and C-terminal (CTD) domains of UreE are in green, teal and pink, respectively. The G1-G5 and CPH motifs of UreG are in cyan, blue, brown, magenta, orange and green, respectively. (E) UreE and UreG forms an asymmetric 2:2 complex, where (F) His68 of distal UreG moves towards the nickel binding site of UreE and chelates a Ni(II) ion with His102 from both proximal and distal UreE. (G-H) Sequence alignment of UreE and UreG. Sequences of *H. pylori* (Hp), *Sporosarcina pasteurii* (Sp) and *Klebsiella aerogenes* (Ka) UreE and UreG were aligned using the program PROMAL3D (30). The names of the secondary structure elements of UreE and UreG follow those used in references (31) and (12). Residue number, according to the Hp sequences, and secondary structure elements are indicated above the alignment. The NTD, CTD, C-terminal tail of UreE, G1-G5 and CPH motifs of UreG are indicated below the alignment.

To avoid the toxic nickel ion escaping into the cytoplasm, nickel ions are delivered within protein complexes. The specificity of nickel transfer is determined by protein-protein interactions – nickel ions are delivered from UreE to UreG and then to urease via the formation of the UreE_2_G_2_ complex and the activation complex (Fig. 1A). Our previous structural studies of protein complexes along the urease maturation pathway – UreFD (14), UreGFD (12), Ni-bound UreG dimer (13) and UreFD/apo-urease complexes (16) – have provided insights into how nickel ions are transferred from UreG to urease via the formation of the activation complex. To shuttle the nickel ions from one protein complex to another, UreG must switch protein binding partners from the UreGFD complex to form the UreE_2_G_2_ complex and then dissociates from the complex after nickel transfer.

The molecular mechanism on how UreE and UreG interact to facilitate urease maturation is not fully understood. Here, we fill the knowledge gap by solving the crystal structures of the UreE_2_G_2_ complex bound with Ni(II) and guanosine 5’-(β,γ-imido) triphosphate (GMPPNP), a non-hydrolysable analogue of GTP. The structures reveal GTP-dependent conformational changes that explain why UreG, in its GTP bound state, prefers to form a 2:2 complex with UreE, and how nickel transfer from UreE to UreG may weaken the UreE-UreG interaction, thereby promoting UreG dissociation from the UreE_2_G_2_ complex. Complemented by mutagenesis and biochemical analyses, we further showed that UreG binding induces conformational changes in the C-terminal tail of UreE, which is important for interacting with the hydrogenase maturation factor HypA but not for interacting with UreG.

## Results

### Crystal structure of UreE_2_G_2_ in complex with Ni(II) and GMPPNP

*H. pylori* UreE contains a C-terminal tail that is not conserved in other species (Fig. 1G). It has been previously shown that truncation of residues 158-170 did not affect the formation of the UreE_2_G_2_ complex (15). To determine the crystal structures of the UreE_2_G_2_ complex, we created two truncation variants of UreE: UreE(1-158) and UreE(1-148) containing residues 1-148 and 1-158 residues, respectively. UreE_2_G_2_ complexes were obtained by mixing the UreE variants with *H. pylori* UreG in 1:1 molar ratio in the presence of Ni(II) ions and GMPPNP, a non-hydrolysable GTP analogue.

The UreE(1-158) complex crystallized in space group P2_1_ with unit cell dimensions of 70.17 × 166.60 × 154.02 Å, α=γ=90°, β= 90.44° (Table S1). Diffraction data were collected to 3.1 Å resolution and the structure was solved by molecular replacement using the crystal structure of UreE (11) as the search template combined with single-wavelength anomalous diffraction (MR-SAD). The UreE(1-158) forms a 2:2 complex with UreG, and four copies of UreE(1-158)_2_G_2_ complexes were found in the asymmetric unit (Fig. 1B). The UreE(1-148)2G2 complex crystallized in space group P2_1_ with unit cell dimensions of 70.97 × 151.07 × 83.15 Å, α=γ=90°, β= 99.50° (Table S1). The structure of UreE(1-148)_2_G_2_ complex was determined to 2.8 Å resolution by molecular replacement using the structure of UreE(1-158)_2_UreG_2_ as the search template. Two copies of UreE(1-148)_2_G_2_ complexes were found in the asymmetric unit (Fig. 1C).

UreE folds into an N-terminal (NTD) and a C-terminal (CTD) domains (Fig. 1D and 1G). The CTD is responsible for dimerization of UreE (Fig. 1E). UreG adopts a SIMIBI-GTPase fold with a seven-stranded β-sheet and ten α-helices (Fig. 1D). The conserved G1-G5 motifs involved in nucleotide binding and GTP-dependent conformational changes are identified in Fig. 1D and 1H.

### UreE forms an asymmetric complex with UreG

In the crystal structure, UreE forms an asymmetric 2:2 complex with UreG (Fig. 1E). UreG forms a dimer that docks to the proximal UreE in a such way that the long axes of UreE and UreG dimers are oriented at an angle of ∼45° (Fig. 2A). The structures of UreE and UreG protomers in the UreE(1-148)_2_G_2_ complex are superimposable to those in the UreE(1-158)_2_G_2_ complex (Fig. S1). In addition, the structures of both UreE protomers are similar, except that the C-terminal residues 148-155 are structured in the proximal UreE but are disordered in the distal UreE (Fig. S1A). Structures of distal UreG only differ from those of proximal UreG in the CPH metal-binding motif (Cys66-Pro67-His68) (Fig. S1B), in which His68 of distal UreG move towards UreE and, together with His102 of UreE, chelates a nickel ion at the UreE/UreG interface (Fig. 1F and S2A). One GMPPNP is bound to each of the proximal and distal UreG and the magnesium ion binding sites of UreG are occupied by the excess nickel ions used in the crystallization (Fig. S2B).

**Figure 2.**
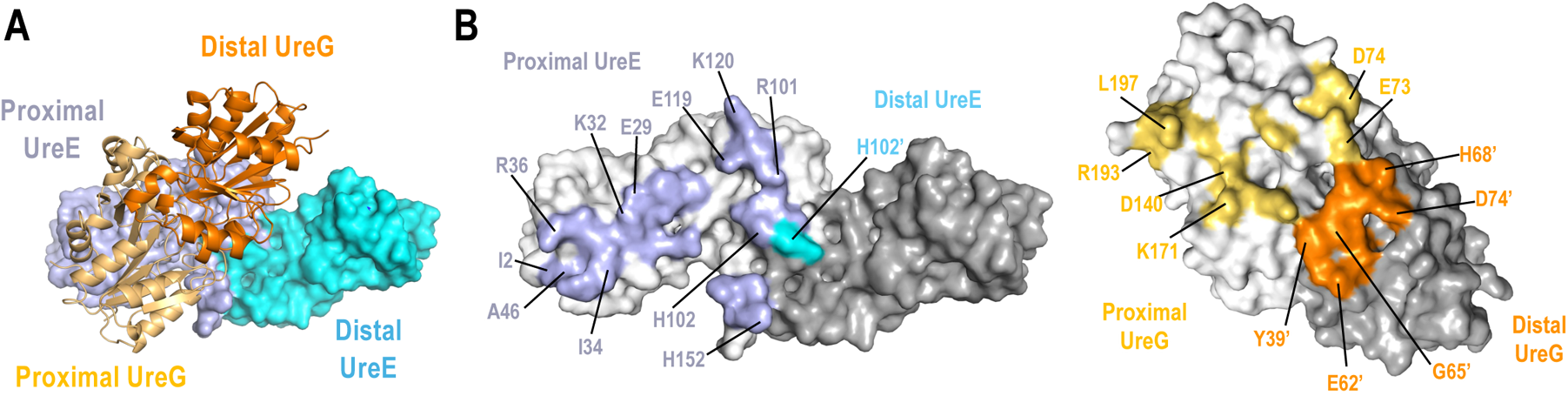
Asymmetric interactions in the UreE₂G₂ complex. (A) UreG dimer (orange) interacts mainly with proximal E (light blue). (B) Residues within 4 Å of the UreE-UreG interface are highlighted in light blue (proximal UreE), cyan (distal UreE), light orange (proximal UreG) or orange (distal UreG). Other residues of distal UreE and UreG are in darker shades of gray. Residues that are directly involved in the UreE-UreG interactions are labeled. Apostrophes indicate residues from distal UreE or UreG.

The interactions between UreE and UreG are asymmetric, with residues of proximal UreE making most of the interactions with UreG (Fig. 2B). Proximal UreE forms a number of hydrogen bonds and salt bridges with proximal and distal UreG (Fig. S3). His102 is the only residue from distal UreE involved in UreE-UreG interactions. This residue, together with His102 of proximal UreE and His68 of distal UreG, chelates a nickel ion at the UreE-UreG interface (Fig. 1F). In addition to polar interactions, Leu197 of proximal UreG docks to a hydrophobic pocket form by Ile2, Ile34, Arg36, Ala46, Arg48 (Fig. S4).

### GTP-dependent conformational changes promote binding of distal UreG to the UreE_2_G_2_ complex

We have previously shown that UreG prefers to form a UreGFD complex in its GDP-bound state, but switches to form a UreE_2_G_2_ complex in the presence of GTP (13). To understand the GTP-dependent conformational changes that promote the formation of the UreE_2_G_2_ complex, we compare structures of proximal and distal UreG to that of the GDP-bound UreG in the UreGFD complex (12) (Fig. 3A). Binding of GMPPNP induces conformational changes in the G2 motif and helix-2 of both proximal and distal UreG. Additional conformational changes are found in the CPH motif in distal UreG (Fig. 3A). GTP-dependent conformational changes are initiated by charge repulsion between the γ-phosphate of GMPPNP and Asp37 of the G2 motif (Fig. 3B). As a result, Asp37 is pushed away from the nucleotide binding site, causing structural rearrangement in the G2 motif and helix-2. In distal UreG, Tyr39 and Gly65 moves towards and form hydrogen bonds with Glu119 of proximal UreE (Fig. 3B). Nickel binding promotes additional conformational changes in the CPH motif of distal UreG – His68 and Asp74 move towards UreE where they interact with the bound nickel ion and Arg101 of proximal UreE, respectively (Fig. 3B). In addition, conformational changes in the G2 motif causes helix-2 to tilt towards the nucleotide binding site, bringing Glu42 and Asp43 of UreG to form inter-subunit salt-bridges with Arg130 and Lys131 of the opposite UreG protomer (Fig. 3C).

**Figure 3.**
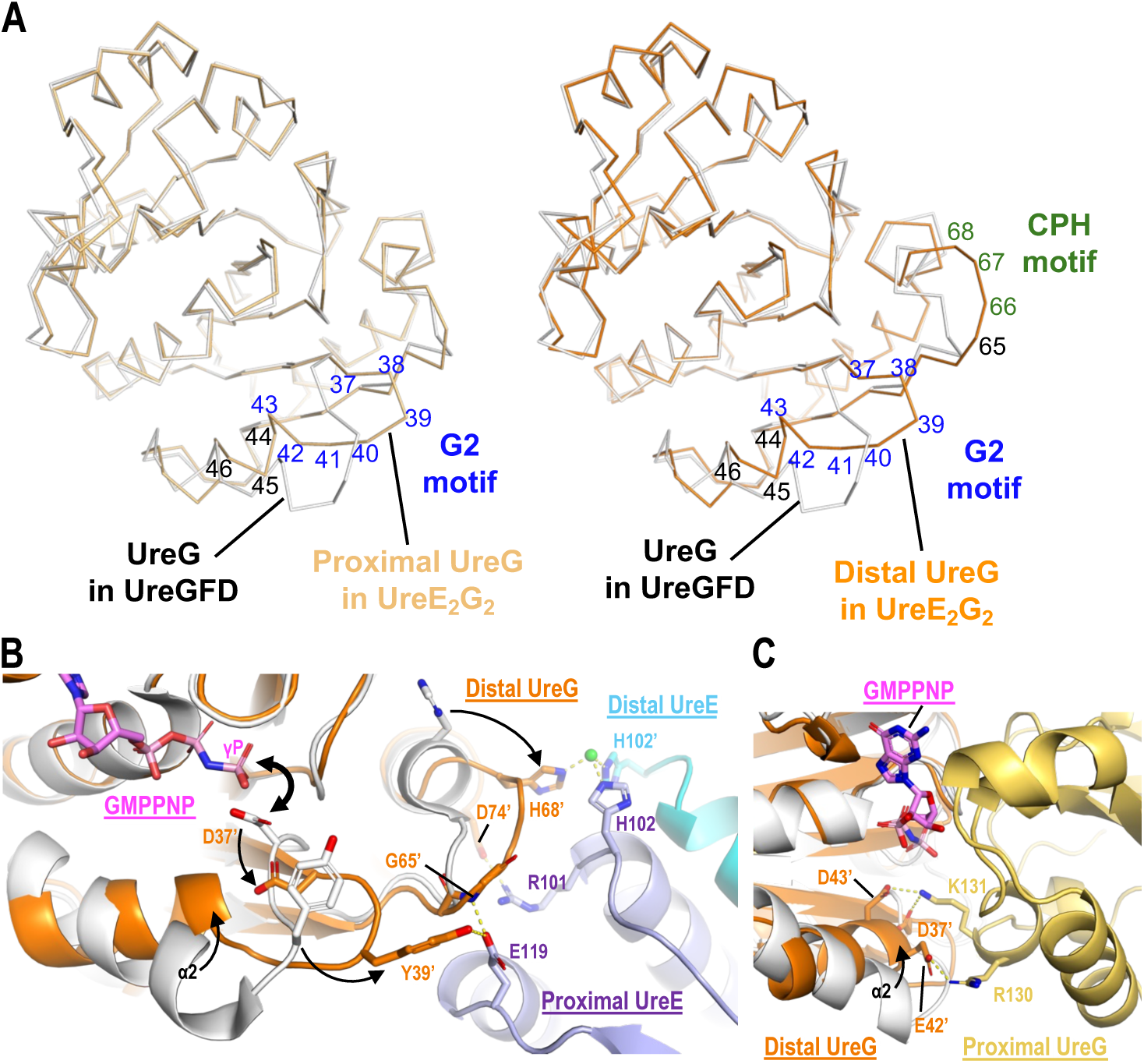
GTP-dependent conformational changes in UreG promote binding of distal UreG to the UreE_2_G_2_ complex. (A) Superimposition of proximal (light orange) and distal (orange) UreG with GDP-bound UreG in the UreGFD complex (gray). GMPPNP binding induces conformational changes in the G2 motif (blue) and helix-2 of both proximal and distal UreG. In the distal UreG, additional conformational changes are found in the CPH metal-binding motif (green). (B) Charge repulsion (thick arrow) between the γ-phosphate of GMPPNP and Asp37 induces conformational changes in the G2 motif. As a result, Tyr39 moves towards and forms a hydrogen bond with Glu119 of proximal UreE. The His68 of the CPH motif of distal UreG moves towards UreE and chelates a nickel ion with His102 from both proximal and distal UreE. Hydrogen bonds are indicated by dotted lines. Apostrophes indicate residues from distal UreE or UreG. (C) Conformational changes in the G2 motif cause helix-2 to tilt towards the nucleotide binding site, bringing Asp37, Glu42 and Asp43 of distal UreG to form salt-bridges (dotted lines) to Arg130 and Lys131 of proximal UreG.

We noticed that the G2 motif of proximal UreG is far away from the UreE-UreG interface (Fig. S5). Since the GTP-dependent conformational changes are localized in the G2 motif of proximal UreG (Fig. 3A), we argue that the proximal UreG in its GDP-bound state should be able to bind UreE, justifying the observation that UreE forms a 2:1 UreE_2_G complex with UreG in the presence of GDP (15). On the other hand, GTP-dependent conformational changes cause distal UreG to form extra interactions with UreE (Fig. 3B) and with proximal UreG (Fig. 3C), explaining why GTP binding promotes the anchorage of distal UreG to form the UreE_2_G_2_ complex.

### Formation of UreE_2_G_2_ complex induces conformational changes in the NTD and C-terminal tail of proximal UreE

The structures of proximal and distal UreE in the UreE_2_G_2_ complex were compared to the crystal structure of the Ni-bound UreE dimer (11). Structural superimposition reveals conformational changes in the NTD and C-terminal tail of proximal UreE (Fig. 4A). As shown earlier, Leu197 of proximal UreG forms hydrophobic contacts with a pocket composed of Ile2, Ile34, Arg36, Ala46 and Arg48 on proximal UreE (Fig. S4). Upon formation of the UreE_2_G_2_ complex, the NTD of proximal UreE undergoes a domain movement that positions these residues to interact with Leu197 of proximal UreG (Fig. 4B).

**Figure 4.**
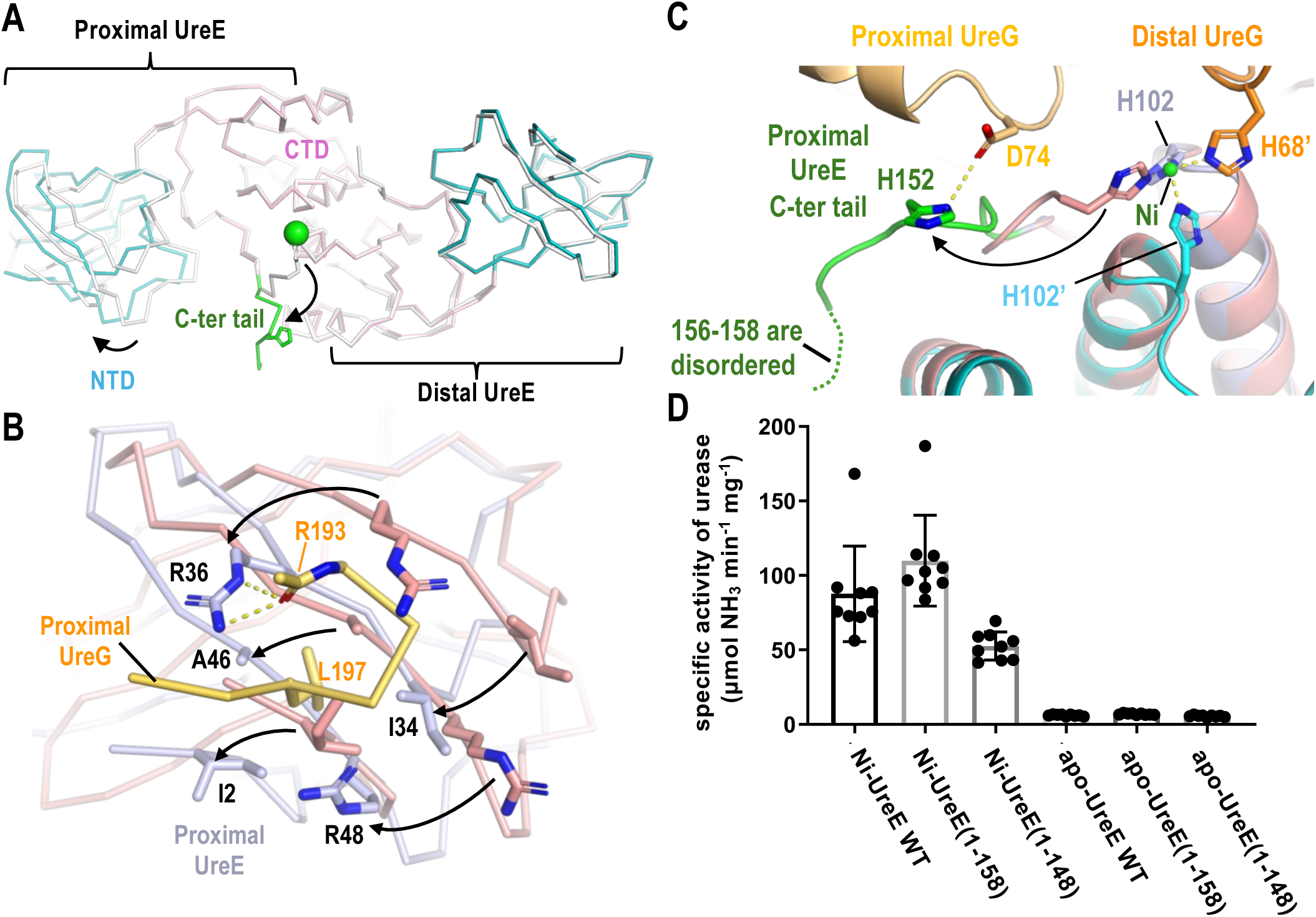
UreG binding induces conformational changes in NTD and the C-terminal tail of proximal UreE. (A) Conformational changes in UreE upon the formation of the UreE_2_G_2_ complex. NTD, CTD and the C-terminal tail of UreE are in teal, pink and green, respectively. Compared to the crystal structure of nickel-bound UreE dimer (PDB: 3tj8; gray), conformational changes are found in the NTD and the C-terminal tail (149-155) of proximal UreE. (B) The NTD of proximal UreE (light blue) undergoes a domain movement such that Ile2, Ile34, Arg36, Ala46 and Arg48 form a pocket to accommodate Leu197 of proximal UreG (light orange). Structure of the UreE dimer is in salmon. (C) In the nickel-bound UreE dimer (PDB: 3tj8; salmon), His152 is interacting with a nickel ion at the dimeric interface. Upon the formation of the UreE2G2 complex, the C-terminal tail of proximal UreE (green) swings outwards such that His152 forms a hydrogen bond with Asp74 of proximal UreG. Residues 156-158 of UreE (dotted lines) are disordered and not involved in binding UreG. (D) The C-terminal tail of UreE is not essential for nickel transfer between UreE and UreG. Ni-bound UreE or its truncation variants were added to a reaction mixture containing GTP, KHCO3, UreGFD and apo-urease. Urease was activated with the addition of Ni-bound UreE or its truncation variants, but not their apo-forms.

On the other hand, in the structure of Ni-bound UreE dimer, His152, together with His102 from both protomers, binds a nickel ion at the dimeric interface (11). Binding of UreG in the UreE_2_G_2_ complex causes the C-terminal tail (149–155) of proximal UreE to swing outward such that His152 forms a hydrogen bond with Asp74 of proximal UreG (Fig. 4C).

To further assess the role of the C-terminal residues of UreE in nickel delivery, we have prepared Ni-bound UreE or its truncation variants and test if they can activate urease in the presence of UreGFD and a reaction buffer containing GTP and KHCO_3_ (Fig. 4D). We also included apo-forms of UreE as negative controls. Our results show that Ni-bound wild-type UreE, UreE(1-158) and UreE(1-148), but not their apo-forms, can activate urease in vitro. There was no significant difference between urease activated by wild-type UreE and UreE(1-158), which is consistent with the observations that residues 156-158 of UreE, which are disordered in the crystal structure of the UreE(1-158)_2_G_2_ complex, are not involved in binding UreG (Fig. S6). On the other hand, urease activated by UreE(1-148) exhibited a lower activity, suggesting that further removal of residue 149-158 would remove the interaction between His152 of proximal UreE and Asp74 of proximal UreG, and thereby weaken UreE-UreG interactions.

### Interaction between HypA and the C-terminal tail of UreE facilitates nickel transfer between the hydrogenase and urease maturation pathway

How UreE receives its nickel ions from upstream sources is not fully understood. In *Helicobacter* species, which contain both the [NiFe]-hydrogenase and urease, there are accumulating evidence suggesting that UreE can receive its nickel ions by cross-talking with the hydrogenase maturation pathway (5). Nickel delivery to the [NiFe]-hydrogenase is facilitated by the hydrogenase maturation factors HypA and HypB. UreE can form a 2:1 complex with HypA (15, 17, 18).

To investigate whether UreE can receive nickel ions by protein-protein interaction with HypA, we performed an in vitro urease activation assay using a two-chamber dialyzer. To simplify protein purification, we attached a Strep-tag II sequence (WSHPQFEK) to the C-terminus of HypA (HypA_str_). We show that urease was activated only when Ni-bound HypA_str_ was mixed with UreE, UreGFD and apo-urease in the same chamber of the dialyzers (Fig. 5A, reaction 1), but not when Ni-bound HypA_str_ was separated from the rest of the protein components by a dialysis membrane (Fig. 5A, reaction 2). These observations suggest that Ni-bound HypA_str_ can provide the nickel source for urease activation via protein-protein interaction, which is prevented by the dialysis membrane in the reaction 2 of Fig. 5A. Urease activation was greatly reduced when UreE was placed in the left chamber and was separated from the rest of the protein components (Fig. 5A, reaction 3), suggesting that HypA cannot pass its nickel ions to protein components other than UreE. Taken together, our results suggest that UreE receives its nickel ions from the hydrogenase maturation pathway by interacting with HypA.

**Figure 5.**
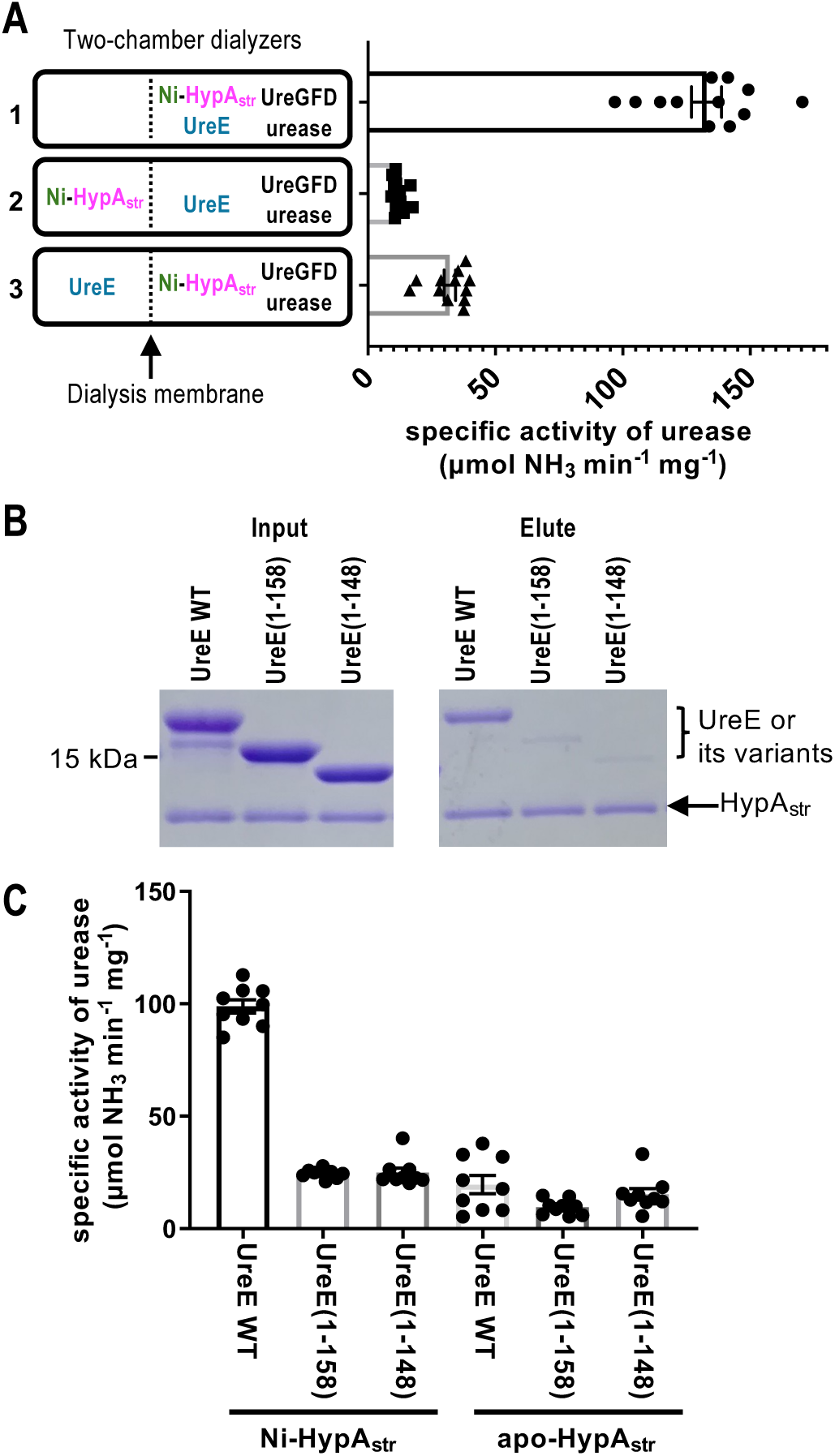
Interaction between HypA and the C-terminal tail of UreE facilitates nickel transfer required for urease maturation. (A) Ni-bound HypAstr (Ni-HypAstr), UreE, UreGFD and apo-urease in buffer containing 2 mM MgSO4 and 300 μM GTP were added to either side of two-chamber dialyzers as indicated. 10 mM KHCO3 was added to activate the GTP hydrolysis of UreG required for urease activation. Urease was activated in A1 where all the protein components for the urease activation were in the same chambers, but not in A2 and A3, where Ni-HypAstr and UreE were separated by a dialysis membrane to prevent protein-protein interactions. Our results suggest that UreE was able to receive nickel from Ni-HypAstr via protein-protein interactions. (B) The C-terminal tail of UreE is essential for HypA-UreE interaction. UreE and its truncation variants were mixed with HypAstr and loaded to Strep-Tactin resins. After extensive washing, bound proteins were eluted with 50 mM biotin. Our results show that truncation of the C-terminal tail of UreE greatly weaken their interactions with HypAstr. (C) Ni-HypAstr was added to UreE or its truncation variants, UreGFD and apo-urease. Apo-form of HypAstr was added as negative controls. Our results suggest that the UreE truncation variants failed to receive nickel ions from HypA for urease activation.

To further examine the interaction between HypA and UreE, we performed pull-down assay by mixing UreE or its truncation variants with Ni-bound HypA_str_ and loaded the mixture to Strep-Tactin resins (Fig. 5B). We showed that wild-type UreE was co-eluted with HypA_str_, suggesting UreE can interact with HypA. In contrast, the interaction between the truncation variants of UreE and HypA_str_ was greatly reduced (Fig. 5B). These results demonstrate that the C-terminal residues 159-170 of UreE is essential for its interaction with HypA.

To access whether the C-terminal residues of UreE is essential for nickel transfer from HypA, we performed an in vitro urease activation assay, where Ni-bound HypA_str_, providing the sole nickel source for urease activation, was added to UreE or its truncation variants, UreGFD and apo-urease. Apo-HypA_str_ was also added as negative controls. Urease activation was observed only when wild-type UreE with an intact C-terminal tail was present (Fig. 5C). Taken together, our results suggest that truncation variants of UreE, lacking the essential C-terminal residues required for interacting with HypA, failed to receive nickel ions from HypA for urease activation (Fig. 5C).

## Discussion

To deliver the toxic Ni(II) ion to the urease, cells have evolved metallochaperones to transport the metal ions from one protein to another. To fulfill their roles in nickel trafficking, metallochaperones must form specific protein complexes like the UreE_2_G_2_ complex to facilitate metal transfer within the complex. They also need to develop a mechanism to dissociate from the complex and switch protein binding partners so that they can shuttle the metal ions to the next protein target along the urease maturation pathway.

The crystal structures of the UreE_2_G_2_ complex reported here show that GTP-dependent conformational changes promote the formation of the UreE_2_G_2_ complex. UreE and UreG form an asymmetric 2:2 complex, in which the proximal UreE makes most of the interactions with the UreG dimer (Fig. 2). GTP binding induces conformational changes in the G2 motif of both proximal and distal UreG that promote the dimerization of UreG (Fig. 3A & 3C). The structural rearrangement in the G2 motif also causes Tyr39 to flip outward and make steric clashes with UreF in the UreGFD complex (Fig. S7) (13), justifying the observation that GTP binding promotes dissociation of UreG from the UreGFD complex (12, 13).

In distal UreG, the conformational changes bring Tyr39 to form a hydrogen bond with Glu119 of proximal UreE (Fig. 3B). In addition, His68 of distal UreG moves towards the nickel binding site of UreE, chelates a nickel ion with His102 of UreE and induces additional conformational changes in the region near the CPH motif of distal UreG (Fig. 3B). As a result, Gly65 and Asp74 of distal UreG form hydrogen bonds with Glu119 and Arg101 of proximal UreE (Fig. 3B). Taken together, GTP binding induces conformational changes in both G2 and CPH motifs, promoting additional interactions between distal UreG and the UreE dimer and the formation of the UreE_2_G_2_ complex (Fig. 3). The role of GTP-dependent conformational changes is also supported by a previous observation that the D37A/E42A variant of UreG, which is defective in GTP-dependent dimerization, failed to switch from the UreGFD complex to form the UreE_2_G_2_ complex (13).

Formation of the UreE_2_G_2_ complex facilitates nickel transfer by juxtaposition of the nickel binding sites of UreE and UreG (Fig. 6A). We have shown previously that protein-protein interaction between UreE and UreG is required for nickel delivery from UreE to urease (13). Yang and co-workers showed that nickel ions are transferred from UreE to UreG, but not in the reverse direction (15). UreG forms a dimer when bound to a nickel ion and GTP (12, 13). The nickel ion is coordinated in a square planar geometry by Cys66 and His68 of the CPH motif from each of the UreG protomers (13). In the structure of the UreE_2_G_2_ complex, the nickel ion, coordinated by His102 of UreE and His68 of distal UreG, is located at the nickel binding of UreE (Fig. S8). So, the structures reported here represent the structure of the UreE_2_G_2_ complex before the transfer of nickel from UreE to UreG.

**Figure 6.**
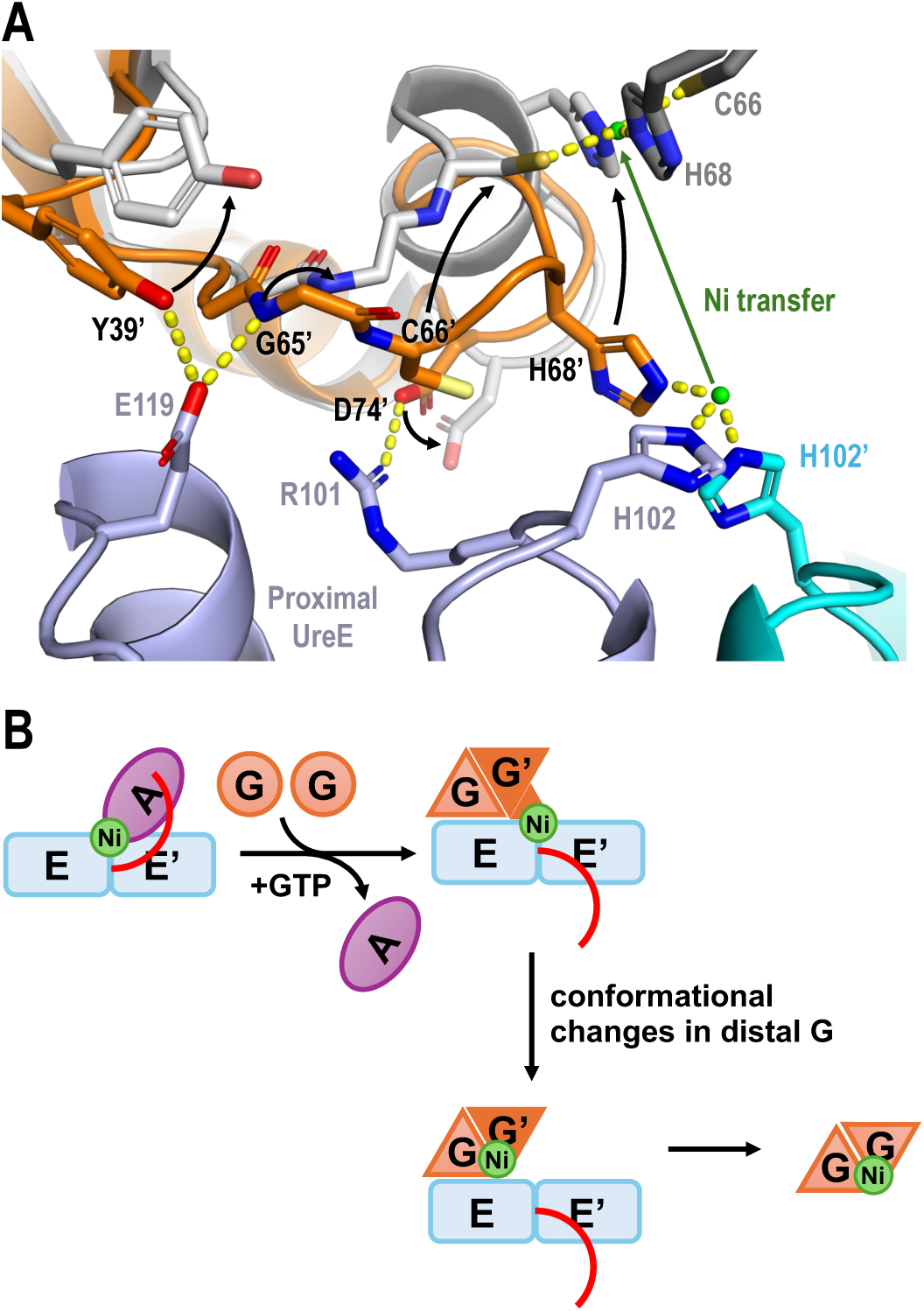
Structural insights into how conformational changes in the UreE_2_G_2_ complex facilitate nickel delivery. (A) Nickel transfer weakens UreE-UreG interactions. The structure of Ni/GMPPNP-bound UreG dimer (gray; PDB: 5xkt) is superimposed with the distal UreG (orange) in the UreE2G2 complex. In the UreE2G2 complex, a nickel ion is bound at the UreE binding site by His68 of proximal UreG and His102 of UreE. Transfer of the nickel ion to UreG (green arrow), where it is coordinated by Cys66 and His68, induces conformational changes (black arrows) that break a number of interactions (dashes) between distal UreG and UreE. Apostrophes indicate residues from distal UreE or UreG. (B) The C-terminal tail (red) of UreE is essential in binding and getting its nickel ion from HypA. GTP-dependent conformational changes promote the binding of the UreG dimer to UreE, which causes the C-terminal tail of UreE to flip outwards and promote the dissociation of HypA. The formation of the UreE2G2 complex facilitates nickel transfer from UreE to UreG, which induces conformational changes in UreG, weakening the UreE-UreG interactions and promoting dissociation of Ni-bound UreG dimer.

To understand the conformational changes in UreG upon transfer of nickel to UreG, we compared the structure of the UreE_2_G_2_ complex to that of the Ni/GMPPNP-bound UreG dimer. Notable differences are found in the CPH motif of distal UreG (Fig. S9) – Cys66 and His68 of distal UreG move back to the nickel binding site of UreG to coordinate the nickel ion at the dimeric interface of UreG (Fig. 6A). Tyr39, Gly65 and Asp74 of distal UreG move away from the UreE-UreG interface, breaking hydrogen bonds with Glu119 and Arg101 of proximal UreE (Fig. 6A). We argue that these conformational changes induced by nickel transfer should weaken the UreE-UreG interaction and facilitate dissociation of UreG.

In *Helicobacter* species, UreE can get its upstream nickel ions by cross-talking with the hydrogenase maturation pathway (5). Pioneer work by Maier’s group shows that knocking out *hypA* and *hypB* genes greatly reduced the activation of both [NiFe]-hydrogenase and urease in *H. pylori* and *H. hepaticus* (19, 20). HypA and UreE can form a HypA-UreE_2_ complex that facilitates nickel transfer between the hydrogenase and urease maturation pathways (15, 17,18). *H. pylori* expressing metal-binding defective mutants of HypA exhibit reduced urease activation and are sensitive to acidic shock (21, 22). All these observations suggest the role of HypA in urease activation in *H. pylori*.

Here, we provided direct evidence that nickel ions from HypA can be delivered to the urease. We showed that urease was activated upon addition of Ni-bound HypA, providing the sole source of nickel ions, to the protein components (UreE, UreGFD and apo-urease) of the urease maturation pathway (Fig. 5A, reaction 1). Separating Ni-bound HypA and UreE by a dialysis membrane in a two-chamber dialyzer greatly reduced urease activation in vitro (Fig. 5A, reactions 2 and 3), suggesting that HypA passes its nickel ion to UreE via protein-protein interactions. We further showed that while removal of the C-terminal residues (159-170) in UreE(1-158) did not affect nickel transfer from UreE to UreG (Fig. 4D), the truncations abolished HypA-UreE interactions (Fig. 5B) and nickel transfer from HypA to UreE (Fig. 5C).

It has been shown that HypA could compete with UreG for binding UreE (23). UreE prefers to form a complex with HypA in the absence of GTP but prefers to form a complex with UreG upon addition of GTP (15). Then what causes UreE to switch protein binding partners? Our work provides structural insights into how GTP-dependent conformational changes promote UreG binding and dissociation of HypA from the HypA/UreE_2_ complex. As discussed, GTP binding induces conformational changes in the G2 and CPH motifs of distal UreG, thereby promotes the anchorage of distal UreG to the UreE_2_G_2_ complex (Fig. 3). We also show that UreG binding induces conformational changes in the residues 149-155 of the proximal UreE, which move away from the nickel binding of UreE (Fig. 4C). Such a movement likely promotes HypA dissociation as the C-terminal residues (159-170) are essential in HypA-UreE interaction (Fig. 5B) (17, 24). In a model by Zambelli and co-workers, the N-terminal MHE motif of HypA was predicted to coordinate a nickel ion at the nickel binding site of UreE (24). As the distal UreG in the UreE_2_G_2_ complex also coordinate a nickel ion at the same location (Fig. S8), the UreG binding is expected to make steric clashes with HypA that promote its dissociation.

A summary on how conformational changes in the UreE_2_G_2_ complex facilitate urease maturation is presented in Figure 6B and Supplemental Movie S1. GTP-dependent conformational changes promote dimerization of UreG and binding of the UreG dimer to UreE. UreG binding induces conformational changes in the C-terminal tail of UreE, which is not essential for UreG binding but essential for HypA interaction, thereby promoting dissociation of HypA. The UreE-UreG interactions are asymmetric, with proximal UreE forming most of the interactions with both proximal and distal UreG. GTP-dependent conformational changes in UreG are also asymmetric – additional conformational changes are observed in the CPH motif of distal UreG, where His68 moves toward and chelates a nickel ion at the binding site of UreE. Nickel transfer from the binding site of UreE to UreG induces conformational changes in distal UreG that weaken the UreE-UreG interactions, and facilitate dissociation of Ni-bound UreG dimer. After dissociation from the UreE_2_G_2_ complex, the UreG dimer then delivers its nickel ion to the urease via the formation of an activation complex with UreFD and apo-urease (16).

Taken together, our work establishes a paradigm on how cells solve the problem of delivering the nickel ion to urease through complexes of metallochaperones. Nickel ions are delivered within the UreE_2_G_2_ complex and the activation complex so that the toxic metal ions do not escape to the cytoplasm. Our work also reveals how GTP and nickel binding allosterically regulate the formation and dissociation of the complexes so that the metallochaperone can shuttle the metal ions to the next protein target along the urease maturation pathway.

## Materials and Methods

### Protein Expression and Purification

The *H. pylori* apo-urease, UreGFD_str_ complex, and UreG were expressed and purified as described previously (13, 16). To prepare protein samples of *H. pylori* UreE and its truncation variants, the coding sequences of wild-type (residues 1-170) UreE, UreE(1-158) and UreE(1-148) were subcloned into the pET-Duet-1 vector to create pHpUreE, pHpUreE(1-158), pHpUreE(1-148), respectively (Fig. S10). These plasmids were then transformed to *E. coli* Rosetta (DE3) pLysS. Bacterial cultures were grown in Terrific Broth with 100 μg/ml ampicillin and 25 μg/ml chloramphenicol to an OD_600_ of 0.6 and induced with 1 mM isopropyl β-D-1-thiogalactopyranoside (IPTG) at 25 °C overnight. Four grams of harvested cells were suspended in buffer SP [20 mM HEPES, pH 6.8, 1 mM Tris(2-carboxyethyl)phosphine hydrochloride (TCEP)] supplemented with 0.1 g of complete ULTRA Tablets protease inhibitor cocktail (Roche) and lysed by sonication. Following centrifugation (20,000 g, 45 min), the supernatant was loaded onto a 5 mL HiTrap SP HP cation exchange column pre-equilibrated with the buffer SP. After washing with 120 mM NaCl in the buffer SP, UreE were eluted using 330 mM NaCl in buffer SP. To remove any bound metal ions in the UreE sample, 1 mM ethylenediaminetetraacetic acid (EDTA) was added to the protein samples, which were then loaded to a HiLoad Superdex 75 PG gel filtration chromatography column pre-equilibrated with the assay buffer (20 mM HEPES, pH 7.5, 200 mM NaCl, 1 mM TCEP).

To prepare protein samples of the UreE_2_G_2_ complex, 150 µM of UreE(1-158) or UreE(1-148) were mixed and 150 µM UreG in the assay buffer supplemented with 2 mM NiSO_4_, 4 mM MgSO_4_, and 5 mM 5’-guanylyl imidodiphosphate (GMPPNP), and purified by loading to a HiLoad Superdex 200 PG gel filtration chromatography column pre-equilibrated in the assay buffer.

To prepare protein samples of HypA_str_, the coding sequences of the Strep-tag II (WSHPQFEK) were added to the C-terminus of *H. pylori* HypA and subcloned into the pRSETA vector to create pHpHypA_str_ (Fig. S10). The plasmid was transformed into *E. coli* Rosetta (DE3) pLysS. Bacterial cultures were grown in Terrific Broth with 100 μg/ml ampicillin and 25 μg/ml chloramphenicol to an OD_600_ of 0.6 and induced with 1 mM IPTG at 25 °C overnight. Three grams of harvested cells were suspended in the Strep-binding buffer (50 mM HEPES, pH 7.5, 200 mM NaCl, and 1 mM TCEP) supplemented with 0.1 g of complete ULTRA Tablets protease inhibitor cocktail (Roche) and lysed by sonication. After centrifugation at 20,000 g, 45 min, the supernatant was loaded onto a Strep-Tactin XT 4Flow affinity column (IBA Lifesciences) pre-equilibrated with the Strep-binding buffer. HypA_str_ were eluted using 50 mM biotin in strep binding buffer.

To prepare protein samples of Ni(II)-bound UreE and Ni(II)-bound HypA_str_, 1 mM NiSO_4_ was added to 200 µM samples of UreE, UreE(1-158), UreE(1-148) or HypA_str_, which were then loaded to a Superdex 75 10/300 gel filtration chromatography column preequilibrated with the assay buffer to remove the excess NiSO_4_.

### Protein Crystallization and Structure Determination

Two millimolar NiSO_4_, 4 mM MgSO_4_, and 5 mM GMPPNP were added to 170 µM protein samples of UreE(1-158)2G2 or UreE(1-148)_2_G_2_ complex. The protein samples and the reservoir solution (0.2 M sodium malonate dibase monohydrate, 0.1 M bis-tris propane, and 18% PEG3350, pH 6.1) were mixed in a 2:1 volume ratio on a cover slip for hanging drop vapor diffusion at 16 °C. Crystals were cryoprotected by soaking 10% of glycerol and flash frozen in liquid nitrogen. For the UreE(1-148)_2_G_2_ complex, diffraction data was collected at 1.5418 Å using an in-house X-ray generator (Rigaku FRE+) and a RAXIS IV imaging plate detector. For the UreE(1-158)_2_G_2_ complex, crystal diffraction data was collected at the absorption peak of nickel at 1.477 Å on the TPS05A beam line of NSRRC. Diffraction data were indexed and integrated using the program XDS (25) and scaled with AIMLESS in the CCP4 suite (26). For the UreE(1-158)_2_G_2_ complex, phases were determined by MR-SAD using the crystal structure of Ni-bound UreE (11) (PDB code: 3tj8) as the search template. For the UreE(1-148)_2_G_2_ complex, phases were determined by the molecular replacement method using the structure of the UreE(1-158)_2_G_2_ complex as the search template. Models were built interactively using the program COOT (27) and refined using the programe PHENIX.REFINE (28). Figures of protein structures were created using PyMOL (www.pymol.org).

### Pull-down assay

For the pull-down assay in Fig. 5B, 150 µM of UreE or its truncation variants and 150 µM of HypA_str_ were added to 0.1 mL Strep-Tactin XT 4Flow resin pre-equilibrated with the Strep-binding buffer in a spin column. After washing with 0.4 mL Strep-binding buffer, proteins were eluted with 50 mM biotin in the Strep-binding buffer, analyzed by SDS-PAGE, and stained with Coomassie Blue.

### In-vitro urease activation assay

We reconstructed the urease maturation pathway in vitro using purified protein samples (Fig. S11). For the urease activation assay in Fig. 4B, 40 µM of Ni-bound UreE, UreE(1-158) or UreE(1-148) were added to 10 µM of apo-urease, 40 µM of UreGFD_str_ in the reaction buffer (20 mM HEPES, pH 7.5, 200 mM NaCl, 1 mM TCEP, 2 mM MgSO_4_ and 300 μM GTP). For the urease activation assay in Fig. 5C, 40 µM Ni-bound HypA_str_ was added to 10 µM of urease, 40 µM of UreGFD_str_, 40 µM of UreE, UreE(1-158) or UreE(1-148) in the reaction buffer. For negative controls, apo-forms of UreE or HypA_str_ were added instead of Ni(II)-bound UreE or Ni(II)-bound HypA_str_. To stimulate GTP hydrolysis of UreG required for urease activation, 10 mM of KHCO_3_ was added to the reaction mixture and incubated at 37°C for 30 min. To measure the urease activity, 50 mM urea was added to the reaction mixture and incubated at 37°C for 30 min and the amount of ammonia released was measured using a phenol/hypochlorite reaction (29).

### Urease activation assay in a two-chamber dialyzer

20 μM Ni(II)-bound HypA_str_, 40 μM apo-form of UreE, 40 μM UreGFD_str_, and 10 μM apo-urease in the reaction buffer were added to either side of two-chamber dialyzers (Bioprobes Ltd.) separated by a dialysis membrane (3.5 kDa MWCO; Spectrum Labs) as indicated in Fig. 5A. The dialyzers were incubated at 37°C for 60 min with shaking. To stimulate GTP hydrolysis of UreG required for urease activation, 10 mM of KHCO_3_ was added to the reaction mixture and incubated at 37°C for 30 min. To measure the urease activity, 50 mM urea was added to the reaction mixture from the urease-containing chamber and incubated at 37°C for 20 min. The amount of ammonia released was quantified using a phenol/hypochlorite reaction (Weatherburn 1967).

## Acknowledgement

This work was supported (in part) by grants from the Research Grants Council of Hong Kong (14117321, 14117223, C4041-18E, C4002-21-EF, C2003-22W, AoE/M-403/16, AoE/M-05/12, AoE/M-402/25-N) and by direct grants from the Chinese University of Hong Kong. Molecular graphics and analyses were performed using PyMOL (https://pymol.org/). We thank the National Synchrotron Radiation Research Center for providing beamline for X-ray data collection.

## Data deposition

The atomic coordinates and structure factors of the UreE(1-158)2G2 and UreE(1-148)_2_G_2_ complexes (PDB ID code: 20ZY and 21AD, respectively) were deposited in the Protein Data Bank, www.wwpdb.org.

---

**Table S1.** Data collection and refinement statistics.

|  | **UreE(1-158)<sub>2</sub>UreG<sub>2</sub>** | **UreE(1-148)<sub>2</sub>UreG<sub>2</sub>** |
| --- | --- | --- |
| PDB code | 20ZY | 21AD |
| **Data collection** |  |  |
| Wavelength (Å) | 1.477 | 1.542 |
| Space group | P2<sub>1</sub> | P2<sub>1</sub> |
| Cell dimensions |  |  |
| a, b, c (Å) | 70.17, 166.60, 154.02 | 70.97, 151.07, 83.15 |
| α, β, γ (°) | 90.00, 90.44, 90.00 | 90.00, 99.50, 90.00 |
| Resolution (Å) | 49.69 – 3.10 (3.18 – 3.10) | 46.93 – 2.85 (2.97 – 2.85) |
| R<sub>merge</sub> | 0.089 (0.592) | 0.097 (0.463) |
| CC(1/2) | 0.985 (0.699) | 0.995 (0.812) |
| I/σI | 7.1 (1.3) | 8.4 (2.4) |
| Unique reflection no. | 62886 (4411) | 40155 (4559) |
| Completeness (%) | 98.1 (97.5) | 99.6 (100) |
| Redundancy | 2.5 (2.6) | 3.6 (3.6) |
| **Refinement** |  |  |
| R<sub>work</sub> / R<sub>free</sub> | 0.2089 / 0.2448 | 0.2426 / 0.2719 |
| **No. of atoms** |  |  |
| Protein | 21509 | 10356 |
| Ligand/ion | 268 | 134 |
| Water | 19 | 25 |
| **B-factors (Å<sup>2</sup>)** |  |  |
| Protein | 80.31 | 63.92 |
| Ligand/ion | 90.25 | 48.94 |
| Water | 49.76 | 53.06 |
| **r.m.s. deviations** |  |  |
| bond lengths (Å) | 0.002 | 0.003 |
| bond angles (°) | 0.483 | 0.569 |
| **Ramachandran** |  |  |
| Favored (%) | 98.13 | 98.66 |
| Allowed (%) | 1.87 | 1.34 |
| Outliers (%) | 0.00 | 0.00 |

\*Values in parentheses are for the highest-resolution shell.

**Figure S1.**
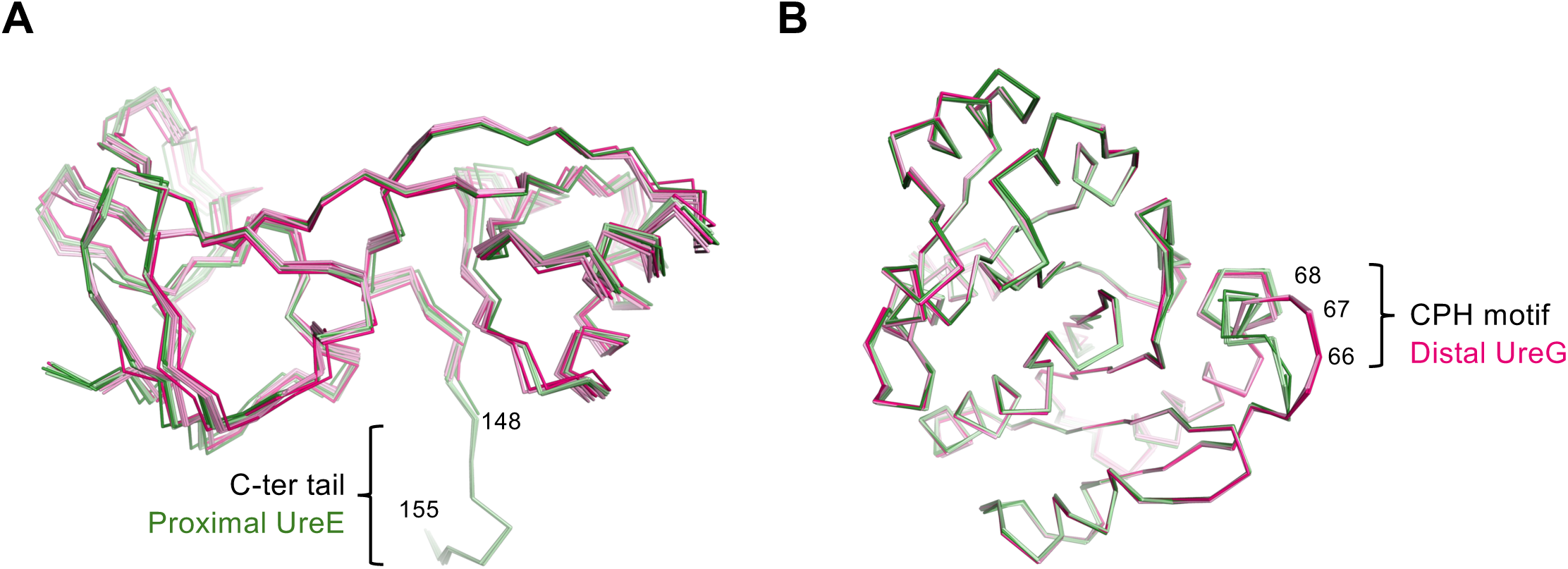
Structural differences between proximal and distal protomers in the UreE_2_G_2_ complex. (A) Structures of proximal UreE (green; chain C, G, K, or O) are superimposable to distal UreE (pink; chain D, H, L, or P). Residues (148-155) in the C-terminal tail are structured in the proximal UreE but are disordered in the distal UreE. The structures of UreE(1-148)_2_UreG_2_ are in darker shades of green and pink. (B) Structures of proximal UreG (green; chain A, E, I, or M) are superimposable to distal UreG (pink; chain B, F, J, or N). The only notable structural differences are found in the metal-binding CPH motif (Cys66-Pro67-His68).

**Figure S2.**
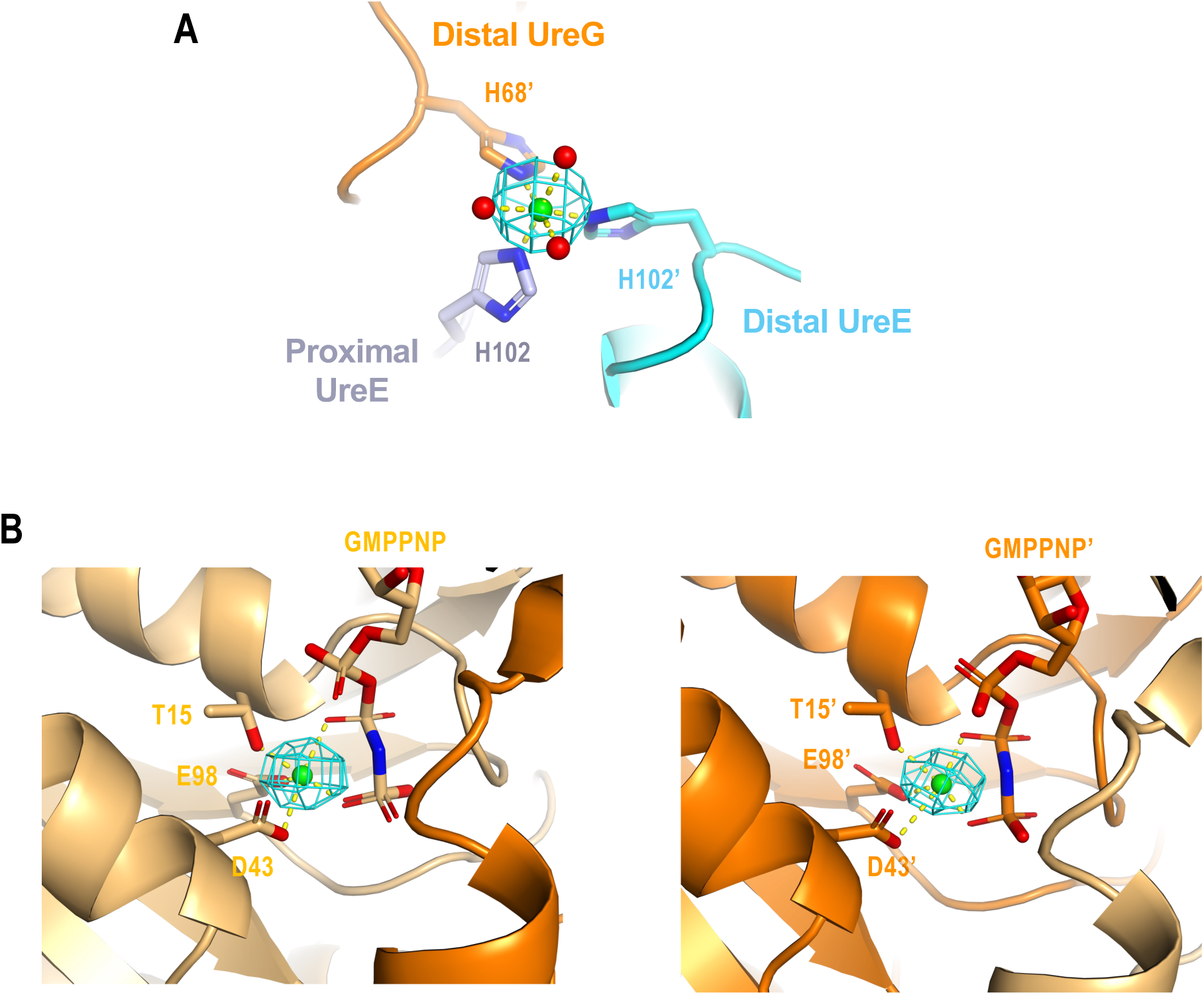
Ni(II) ions identified by anomalous diffraction in the UreE(1-158)_2_-UreG_2_ complex. Anomalous difference map (mesh) was generated by PHENIX using diffraction data collected at the nickel peak wavelength and contoured at 7σ. (A) A Ni(II) ion (green) was chelated by His68 of distal UreG, His102 of proximal and distal UreE and three water molecules (red) in an octahedral geometry. (B) Ni(II) ions occupy the Mg(II) binding sites from each of the nucleotide binding sites of proximal (lef) and distal (right) UreG. Apostrophes indicate residues from distal UreE and UreG.

**Figure S3.**
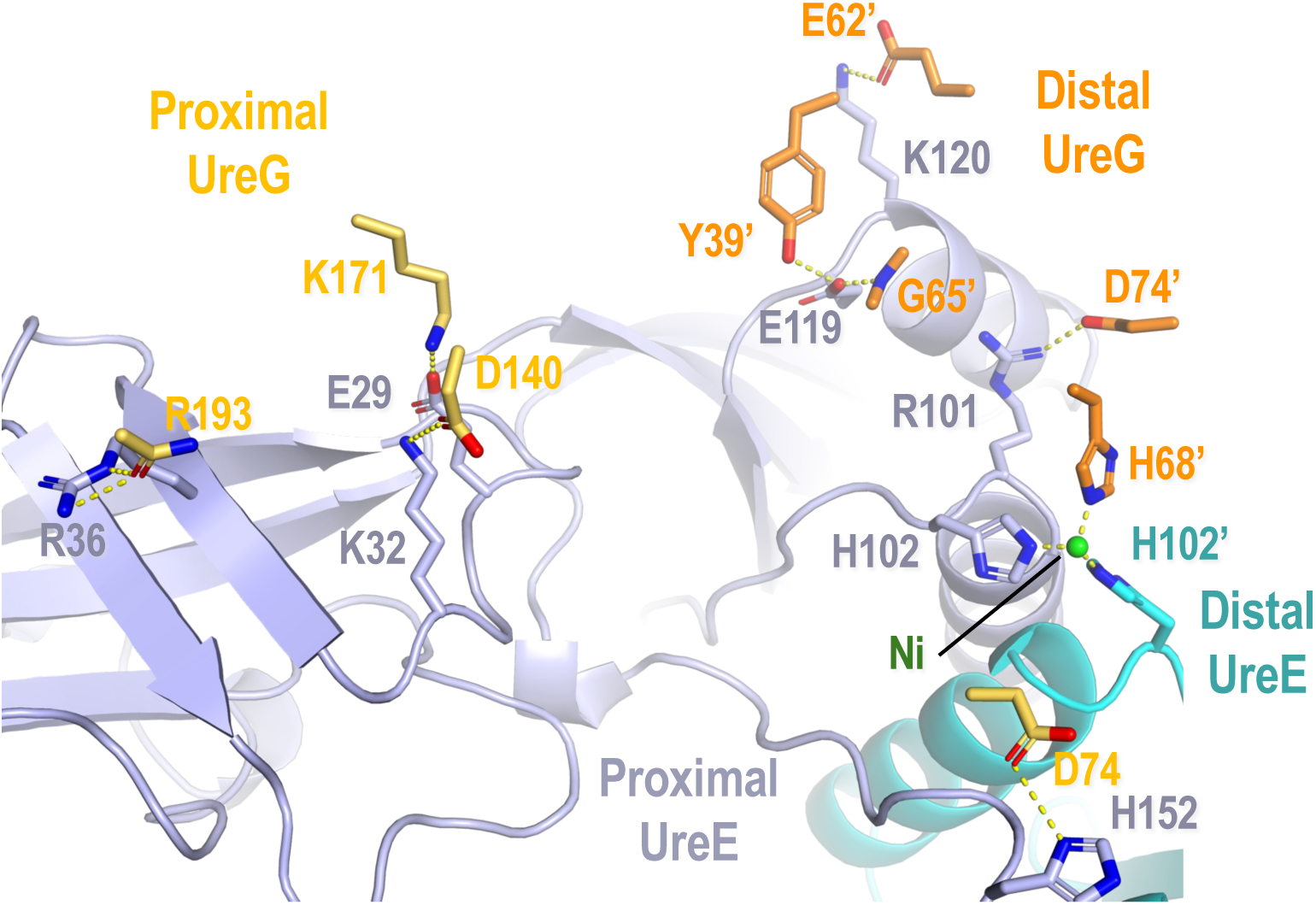
Polar Interactions between UreE and UreG in the UreE_2_G_2_ complex. Residues involved in hydrogen bonds and salt-bridges between UreE and UreG are represented in sticks. Backbone of proximal (light blue) and distal (cyan) UreE are shown as cartoon representation. Residues of proximal and distal UreG are in light orange and orange, respectively. Note that both protomers of UreG mainly interact with proximal UreE. His102’, chelating a Ni(II) ion at the interface with His102 of proximal UreE and His68’ of distal UreG, is the only residue from distal UreE involved in UreE-UreG interaction. Apostrophes indicate residues from distal UreE or UreG.

**Figure S4.**
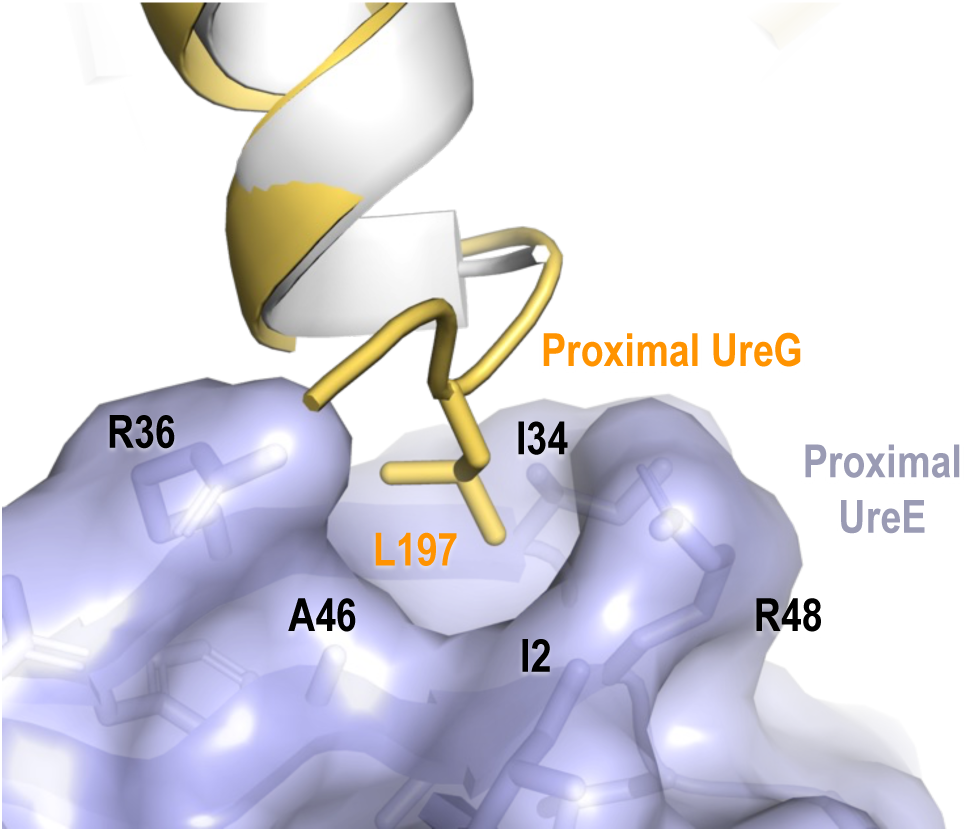
Leu197 of proximal UreG forms hydrophobic interactions proximal UreE. Residue 197-199 of UreG is disordered in the crystal structure of the UreGFD complex (gray) (PDB: 4hi0). These residues become structured in the proximal UreG (light orange) and Leu197 is docked to a hydrophobic pocket formed by Ile2, Ile34, Arg36, Ala46 and Arg48.

**Figure S5.**
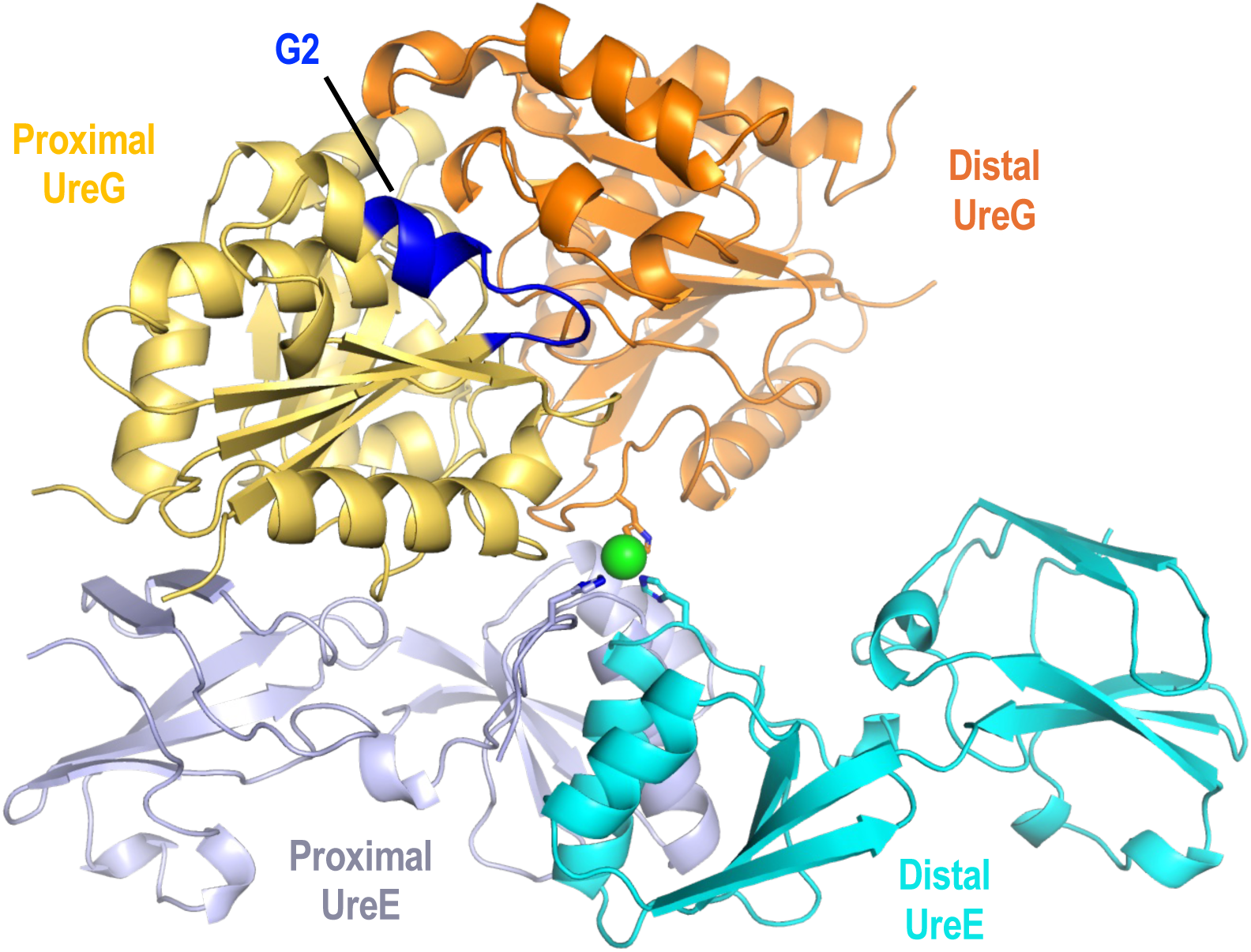
The G2 motif of proximal UreG does not involve in binding UreE. The G2 motif (37–46) of proximal UreG (blue) is located far away from the UreE-UreG interface and does not involve in interacting with UreE.

**Figure S6.**
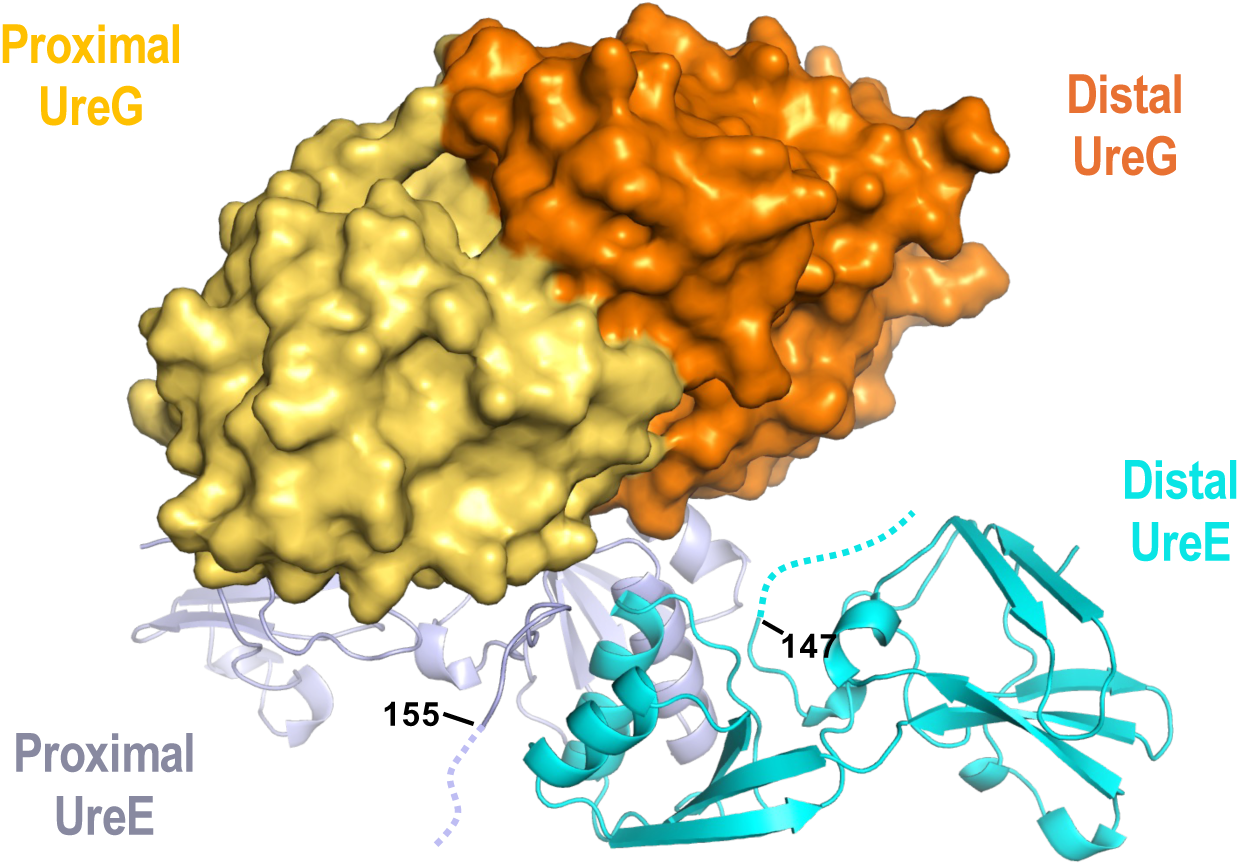
The C-terminal tail of UreE does not involve in binding UreG. In the crystal structure of UreE(1-158)_2_UreG_2_, residues 1-155 of proximal UreE and 1-147 of distal UreE are structured. The rest of the C-terminal tails of UreE are disordered and do not involve in binding UreG.

**Figure S7.**
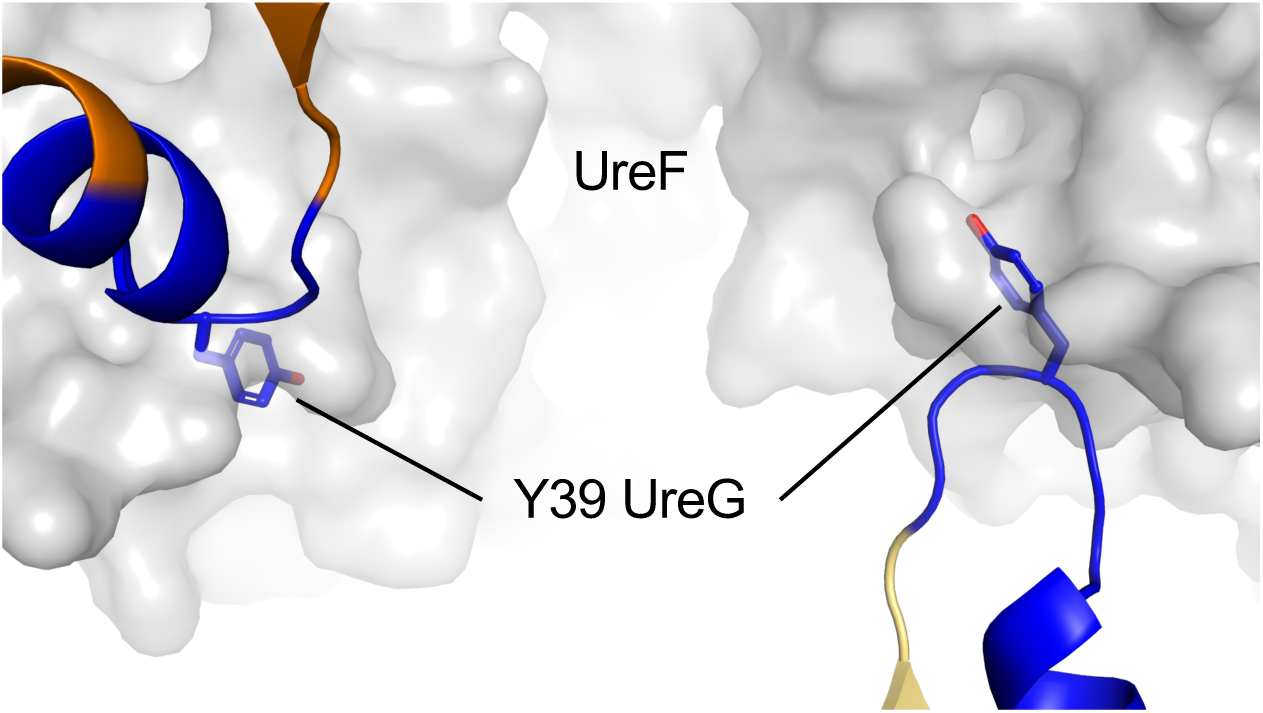
GTP-dependent conformational changes in the G2 motif promotes dissociation of UreGFD complex. GTP binding induces conformational changes in the G2 motif (blue) of UreG such that Tyr39 flips outward and clashes with UreF (gray) in the UreGFD complex. UreF and UreG are in surface and cartoon representations, respectively.

**Figure S8.**
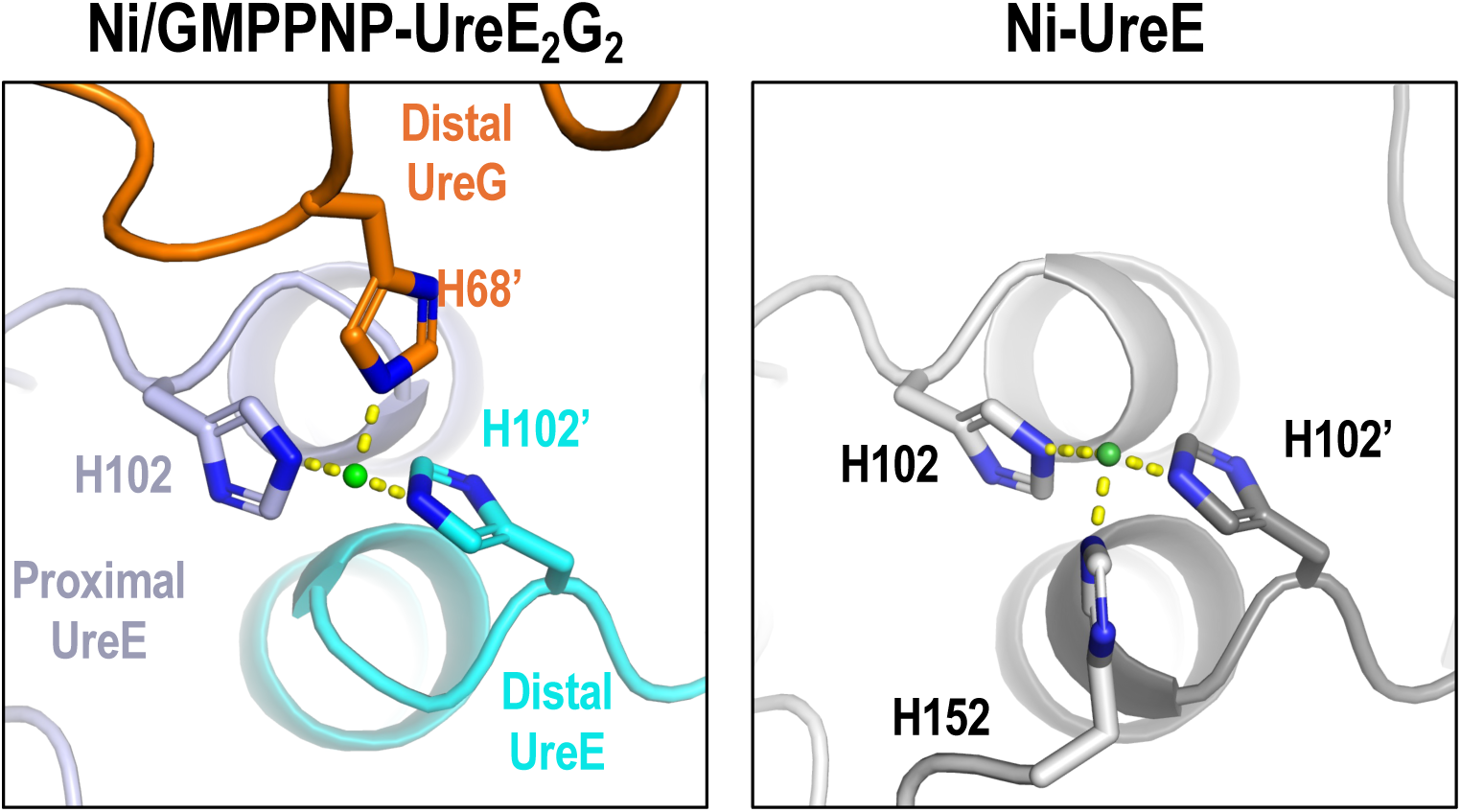
The nickel binding site of the UreE_2_G_2_ complex is the same as that of the UreE. The locations of bound nickel ions in the UreE_2_G_2_ complex (left) and the Ni-bound UreE dimer (right) (PDB: 3tj8) are shown.

**Figure S9.**
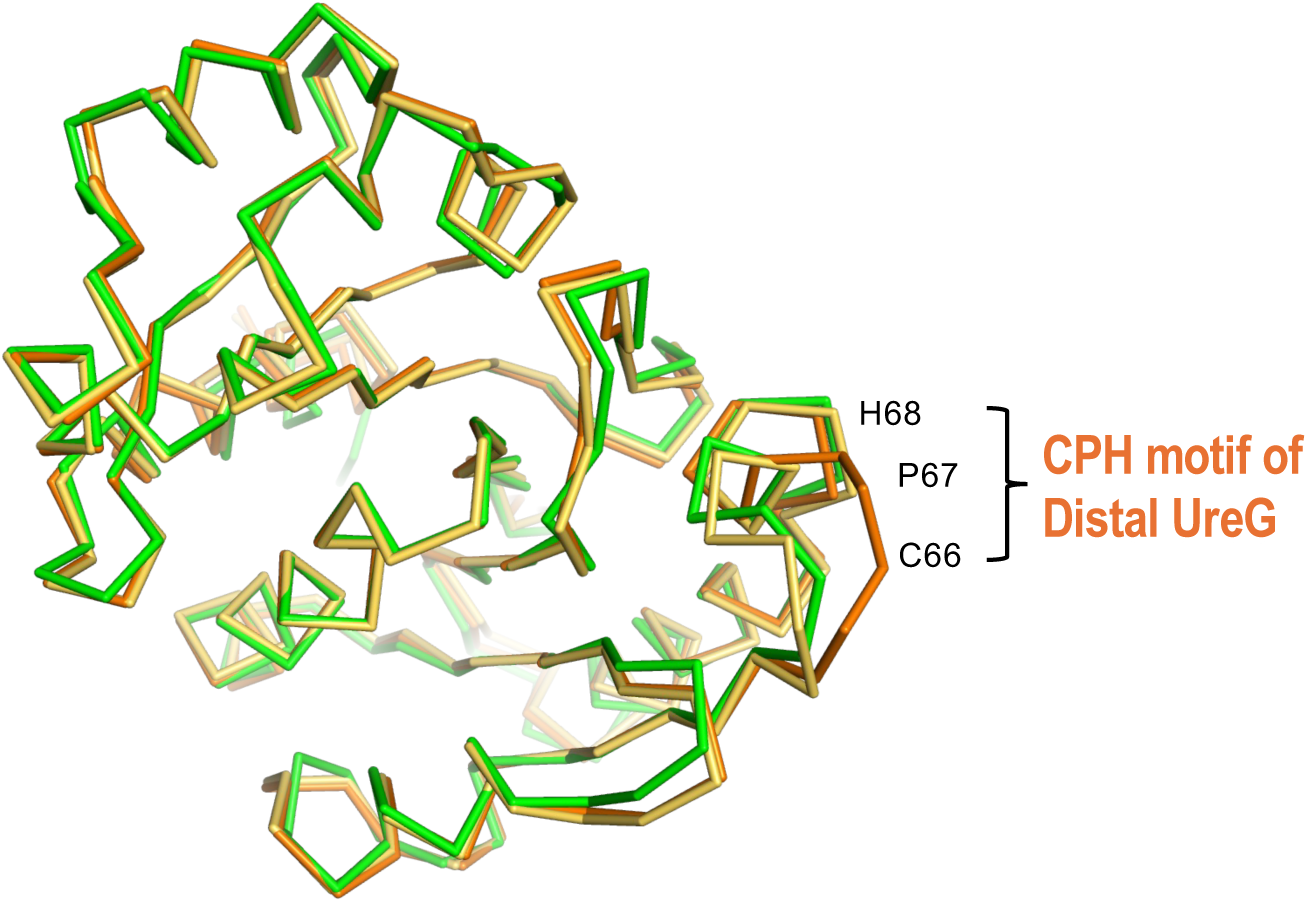
Formation of nickel-bound UreG dimer induce conformational changes in the distal UreG. The structures of proximal (light orange) and distal (orange) UreG were superimposed with Ni/GMPPNP-bound *Klebsiella* UreG (green; PDB: 5xkt). Major conformational changes are found in the CPH motif of the distal UreG.

**Fig. S10.**
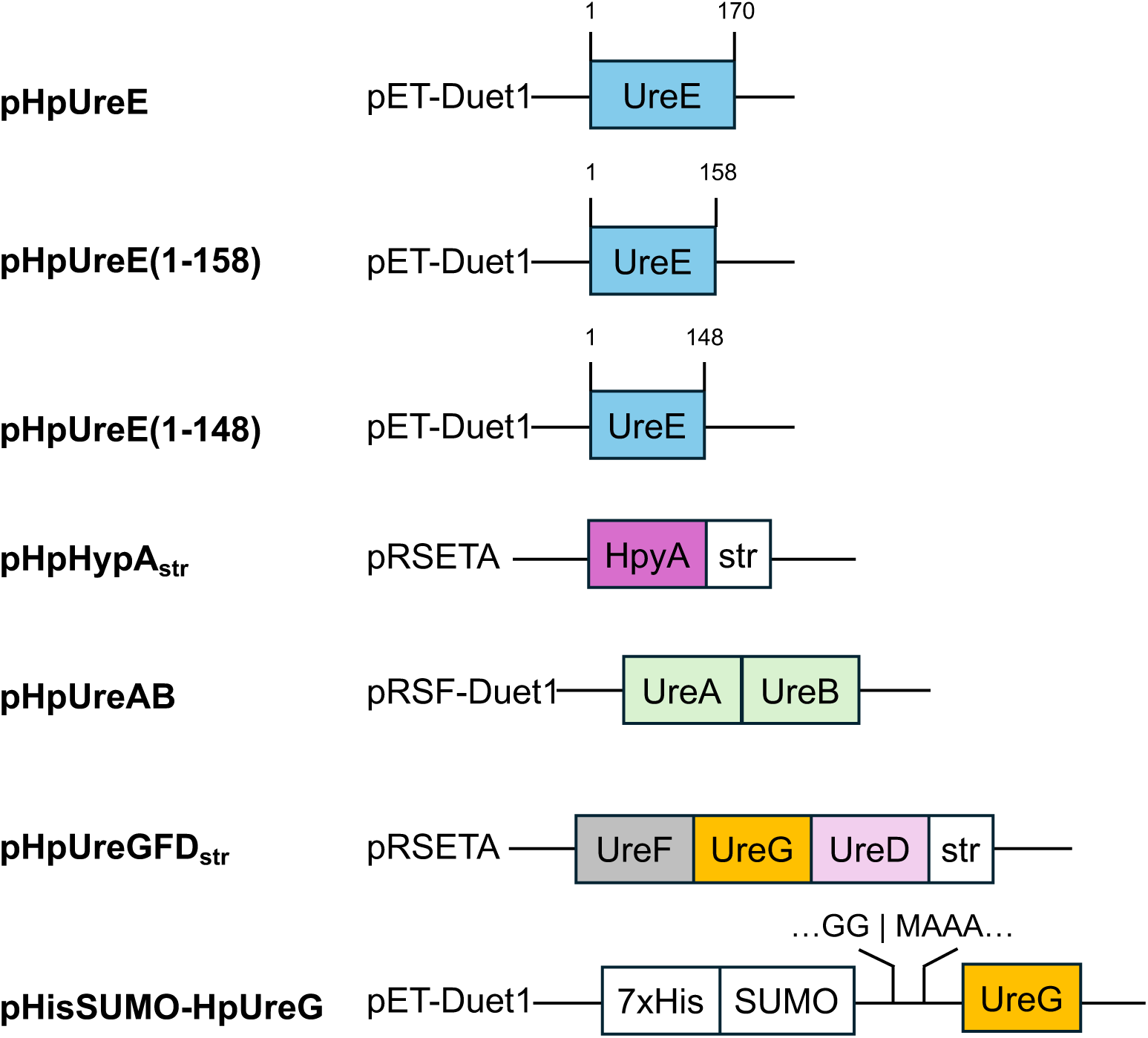
Summary of expression plasmids used in this study. The names of the plasmids are labeled on the left. str: Strep-tag II sequence (WSHPQFEK); SUMO: Small ubiquitin-like modifier protein; 7xHis: poly-histidine tag. The sequences franking the cutting sites of SUMO protease is labeled.

**Figure S11.**
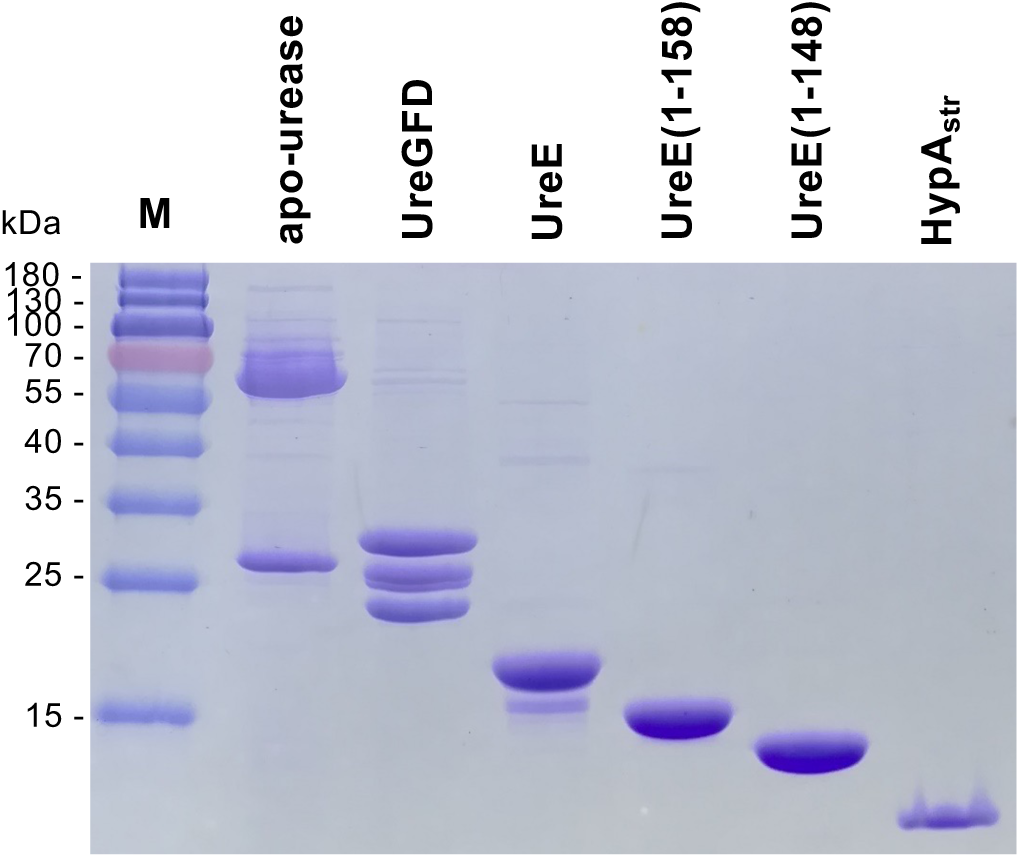
Purified protein samples used in the urease activation and pull-down assays. Purified protein samples of apo-urease, UreGFD, UreE and its truncation variants, HypA_str_ were analyzed by SDS-PAGE stained by Coomassie blue. ThermoFisher PageRuler ™ Prestained Protein Ladder (M) was loaded on the left and the molecular weight of the protein standards were shown.

